# Co-isolation of genetically distinct *Burkholderia pseudomallei* strains from a single patient in North Queensland

**DOI:** 10.1101/2025.09.09.675021

**Authors:** Pauline M.L. Coulon, Piklu Roy Chowdhury, Ian Gassiep, Kay Ramsay, Aven Lee, Edita Ritmejeryte, Miranda E. Pitt, Joyce To, Sarah Reed, Patrick N. A. Harris, Garry S.A. Myers

**Author notes:** Corresponding author: Pauline M.L. Coulon, Australian Institute for Microbiology and Infection, Faculty of Science, University of Technology Sydney, NSW, Australia.

## Abstract

While investigating colony morphology variation (CMV) within clinical isolates of *Burkholderia pseudomallei* (*Bp*) isolated from a single infection, we identified two distinct strains (based on their multi-locus sequence type) from *Bp* TSV292. Simultaneous co-infection is rare, previously reported in only 2 of 133 cases. Here, we present the first comprehensive multi-omics characterization of co-infecting *Bp* strains isolated from a single infection at a hospital in Townsville, Australia in 2018. This finding impacts the design of future diagnosis and treatment of *Bp*.

**Importance:** Melioidosis, a severe disease caused by the bacterium *Burkholderia pseudomallei*, is rarely caused by more than one strain at the same time. In this study, we examined two bacterial strains isolated at once from a patient treated for melioidosis at a hospital in northern Queensland (Australia). By comparing their genomes, proteins, and physical traits, we found important strain differences that may affect how they adapt and survive during infection. Considering strain diversity when studying how this pathogen causes disease is essential, as it directly impacts future diagnosis and treatment.

## Introduction

*Burkholderia pseudomallei* (*Bp*), is an opportunistic pathogen and the causative agent of melioidosis, a severe and often fatal disease in humans and animals (1, 2). Previously endemic to tropical and subtropical regions of Asia, Africa, Central and South America, and northern Australia (1), it is now recognized as established in the United States (3). In most cases, *Bp* acquisition and infection will not lead to the development of melioidosis in healthy populations. In people with underlying conditions, such as diabetes, chronic kidney or lung disease, alcoholism, or compromised immune systems, melioidosis often manifests as skin abscesses or severe pneumonia, which can progress to fatal sepsis with mortality rates reaching up to 52% (4). With climate change intensifying extreme weather events such as heavy rainfall and flooding, the incidence of *Bp* infections is projected to rise (5). In fact, Queensland (Australia), has recently reported its worst year on record for melioidosis, with over 239 confirmed infections and 31 deaths linked to dispersal of the pathogen during flooding events (<u>Queensland Health</u>). While direct transmission between animals and humans has not yet been confirmed, growing concerns have emerged regarding potential transmission from pets, livestock or tropical animals to human (2), either through contact with infected wounds or via shared exposure to contaminated environments (6). *Bp* is naturally highly resistant to several antibiotic classes. Resistance to antibiotics used as primary treatment (meropenem, ceftazidime and trimethoprim/sulfamethoxazole) is uncommon (7), however emerging resistance against these drugs is increasingly described (8–11).

A key yet often overlooked feature of *Burkholderia* species is their ability to undergo colony morphotype variation (CMV), a reversible process by which bacteria switch between distinct colony phenotypes (12). This phenomenon reflects rapid adaptation to environmental stressors and is often associated with changes in outer membrane proteins, antibiotic production, and virulence factors (13–17). Mechanisms underlying CMV often include mutations in global regulators, two-component systems, genome reduction and duplication, bacteriophage integration, and DNA methylation (recently reviewed by Coulon et al. (12)). *Bp* is known to exhibit up to seven different macrocolony morphologies which can be modulated under laboratory conditions (13, 17, 18). Importantly, CMV is not limited to adaptation *in vitro* as it also occurs during infection (19). Approximatively 10% of 450 *Bp* clinical isolates produce mixed mucoid and non-mucoid CMV populations on blood agar (19). In a long-term murine infection model, small colony variants (scv) were observed (20, 21). During human infection, differences in O-antigen lipopolysaccharides (LPS; [OPS]) production explains the mucoid phenotype (as observed on blood agar) even in the absence of mutations or altered expression of the *wbiA* O-antigen acetylase (19). Additionally, genes within the LPS biosynthesis cluster are upregulated during persistent infection in mice (20, 21). The upregulation of the σ-54 dependent regulator *yelR*, is involved in the emergence of two reversible yellow *Bp* variants, which display enhanced resistance to hypoxic stress and increased colonization and persistence in the murine stomach (22).

While investigating CMV within a cohort of *Bp* isolates (23) collected from clinical melioidosis cases presented at Townsville University Hospital (Australia) (24), we identified a polyclonal infection initially mistaken for colony morphotype variants. The apparent rough and smooth ‘variants’ were in fact two genetically distinct strains isolated from the same infection site. Polyclonal *Bp* infection is rarely reported, with an apparent occurrence of 1.5% in a cohort of 133 patients (25); to date no integrated multi-omics analyses have been performed at the bacterial level. Here, we present the first study combining genomic, proteomic, and phenotypic analyses of two co-infecting *Bp* strains from a melioidosis patient in Australia.

## Materials and Methods

### Ethical Approval

This study received ethical approval from the Royal Brisbane and Women’s Hospital Ethics Committee (LNR/2020/QRBW/65573), with site-specific authority obtained from the Townsville Hospital and Health Service and approval under the Queensland Public Health Act.

### Colony morphology screening

*Bp* TSV292 was streaked on LB agar plates containing 0.01% Congo red (CRLA) from - 80°C stock and incubated at 37°C for two overnight periods.

### Genomic analysis

#### gDNA extraction

Using either gDNA spin or high molecular weight gDNA extraction kits (NEB), gDNA was extracted from multiple macrocolonies grown for three overnights at 37°C on CRLA. Short-read sequencing (Illumina) was conducted on all CMVs. Briefly, libraries were prepared using 1 ng of gDNA via the Nextera XT kit and sequenced using the NovaSeq XPlus platform (35). On select isolates, long-read sequencing (Oxford Nanopore Technologies) was implemented. Libraries were prepared using 400 ng gDNA with the SQK-NBD114.24 kit and sequenced using R10.4.1 flow cells on the GridION platform. Reads were base called using Dorado 7.6.7.

#### Genome assembly

Nanopore and Illumina reads were filtered and trimmed using respectively nanofilt (v.2.8.0; (26)) and fastp (v0.24.1; (27)). Each set of Nanopore long reads were then assemble using autocycler tool (v0.4.0; (28)). Next, the resulting long-read consensus assemblies were polished with Illumina short reads using polypolish (v0.6.0; (29)) and then pypolca (v3.0.1; (30)). Finally, the obtained assembly was annotated using batka (v1.11; (31)). For the second CMV isolated from each isolate, Illumina short reads were assembled using unicycler short read assembler (v.0.5.1; (32)) generating contigs.

#### Genomic modification analyses

Each set of Illumina reads, belonging to the second CMV, were mapped on their respective first CMV assembly using snippy tools with haploid settings (v4.6.0; (33) available on Galaxy) to determine SNPs and indels, to confirm the results, the first CMV Illumina reads were also mapped onto their own assembly, resulting in showing no genomic variation.

#### Genotyping and Phylogeny analysis

6,259 genomic allelic profiles of 1,225 available *Bp* isolates on pubMLST were retrieved and used to generate a distance phylogenetic tree with GrapeTree using allelic profile differences (34). The resulting tree was generated using ggtree R package. <u>PhyloSift</u> v1.0.1 was used to resolve subclade phylogeny of the two clusters (identified from the wgMLST clustering analysis) by computing a maximum-likelihood tree based on the genomes (available on pubMLST) using FastTree2.2 (35). <u>FigTree</u> v1.4.4 was used to draw the unrooted phylogenetic trees and <u>FastANI</u> was used to identify the pairwise average nucleotide identity with the respective ST identified in this analysis. BRIG-0.95 (36) was used to map pairwise BLASTn results of 5 to 6 six genomes (37)-selected from the PhyloSift phylogeny.

#### Prediction of antibiotic resistance genes

AMR gene orthologs present in genomes was identified using the CARD database (38) and outputs were filtered to tabulate data of candidates which had 100% coverage of subjects. Antibiotic Resistance Detection and Prediction (ARDaP) tool was used to determined antibiotic resistance profile of both isolates (39).

#### Whole genome alignment

Both MAUVE (40) and LASTZ (41, 42) tools were used to aligned TSV292_1 (rough) and TSV292_2 (smooth) together, using the default settings. To correct aligned genomes with LASTZ tool, the first residue numbers for each chromosome were changed as followed, to set up the same origin. For TSV292_1 rough contig 2 residue 1,249,185 became 1 while TSV292_2 (smooth) contig 1 was reversed and residue 285,486 became 1; TSV292_2 (smooth) contig 2 residue 490,391 became 1.

### Methylome analysis

#### Identification of genes encoding for DNA methyltransferase and their linked motifs

Pod5 files obtained from Nanopore sequencing were merged, using pod5 merge function and analyzed for DNA methylation motifs using the Dorado with both <u>dna_r10.4.1_e8.2_400bps_hac@v4.3.0_6mA@v2</u>, dna_r10.4.1_e8.2_400bps_hac@v4.3.0_5mC_5hmC@v1 models (https://github.com/nanoporetech/dorado ). To identify the location and number of detected motifs, as well as the genes encoding DNA methyltransferases, the call_methylation and annotate_rm functions from MicrobeMod (43) were used. These tools were provided with the corresponding genome assembly “.fasta” and annotation “.gff3” files for each isolate.

### Proteomic analysis

#### Protein extraction and peptide preparation

Bacterial cells in 1M Tris-HCL containing 1% SDS were sonicated in an ultrasonic bath YJ5120-1 (Labtex) for 3 minutes, followed shaking at 1,000 rpm at 95°C for 15 minutes on a Thermomixer Compact (Eppendorf). The mixture was allowed to cool down to room temperature and 30 µL was mixed with 270 µL of Buffer I (6 M guanidine chloride, 50 mM Tris pH 8.0, 10 mM DTT) and incubated at 30°C for 40 min. Following addition of 9 µL of 0.5 M iodoacetamide, the sample was incubated at 30°C for 50 min. The reduced and alkylated protein sample was transferred to a 10 kDa cut-off Amicon Ultra device (Merck) which was placed into a collection tube and centrifuged at 14,000 × g for 30 min. The flow-through from the column was removed and 300 µL of 100 mM ammonium bicarbonate was added to the column followed by centrifugation at 14,000 × g for 30 min. To digest proteins in the column, 130 µl of 100 mM ammonium bicarbonate containing 2 µg Trypsin was added. The column was incubated at 37°C for 24 h and then centrifuged in a new collection tube at 14,000 × g for 30 min. Fifty µL of water with 0.1% formic acid was then added to the spin column followed by centrifugation at 14,000 × g for 30 min to elute the remaining digested peptides. Enzymes and salts were removed using C_18_ ZipTips (Merck) following the manufacturer’s instructions and desalted peptides were eluted with 80% acetonitrile/0.1% formic acid into microcentrifuge tubes. Digested peptide samples were dried down using a vacuum concentrator (Concentrator plus, Eppendorf) and reconstituted with 50 μL water with 0.1% formic acid prior to LC-MS/MS analysis.

#### LC-MS/MS analysis

The samples were chromatographically separated and resulting proteomics data was acquired using an Acquity M-class micro-LC system (Waters, USA) coupled with ZenoTOF 7600 LC-MS/MS system using the OptiFlow Turbo V ion source. (AB Sciex). Two µL of sample were loaded onto Micro TRAP C18 column (0.3 × 10 mm; Phenomenex) and washed for 10 min at 10 µL/min with 97% solvent A (water with 0.1% [v/v] formic acid) and 3% solvent B (99.9% [v/v] acetonitrile, 0.1% [v/v] formic acid). Liquid chromatography was performed at 5 µL/min using a 2.7 µm Peptide C18 160Å; (0.3 × 150 mm; Bioshell) column with the column oven temperature maintained at 40°C. The gradient started at 3% solvent B which was increased to 5% by 0.5 min, followed by an increase to 35% solvent B by 38.9 min and then another increase to 80% solvent B by 39 min. The column was washed with 80% solvent B for 3.9 min before equilibrating with 3% solvent B from 43 to 45 min. Data was acquired using Zeno SWATH DIA, using a 65 variable width SWATH windows. The TOF MS parameters were as follows: precursor mass range was 400 – 1500 m/z; declustering potential (DP) was set to 80 V and accumulation time was set at 0.25 s. The TOF MS/MS parameters were as follows: fragmentation mass range was set to 100 – 1500 m/z; with an accumulation time of 0.025s; fragmentation mode was set to CID with Zeno pulsing selected. The collision energy for the MS/MS acquisition was automatically adjusted according to the m/z and charge of the peptide.

#### Data analyses

Raw label free quantification (LFQ) LC-MS/MS data were converted to .mzML files, using MS convert, to allow the data search using Fragpipe v22 (44). Protein identification was conducted against the respective annotated proteomes using the DIA_SpecLib_Quant workflow, which builds a spectral library with MSFragger-DIA and performs quantification with DIA-NN. The output “diann-output.pg_matrix.tsv” was filtered to remove contaminants and retain proteins with at least two non-missing values for at least one group. Data was normalized using the variance stabilization normalization (45). Proteins with two or more missing values in one group were categorized as “detected/not detected.” For proteins with a single missing value in one group, missing data were imputed using the mean of the other replicates in that group. Differential protein expression between groups was assessed using Welch’s t-test, with Benjamini–Hochberg correction applied to control the false discovery rate (adjusted p < 0.05) (46, 47). Analyses were performed on three biological replicates (Table S5-9).

#### Proteomic representation

To assign homologous proteins between TSV292 rough, smooth and K96243 (*Bp* model), a megablastp analysis was set up with only one output match for each gene in query using the following parameters: -evalue 10 -outfmt 6 -num_threads 3 - max_target_seqs 1 -max_hsps 1 (37). Clustering of orthologous groups (COGs) for annotated proteins were determined using EggNOG tool v5 ( (48)). To represent the data in a volcano plot, detected/not detected data were imputed using the lower normalized Log2 value for intensity with an imputed adjusted pvalue randomly assigned between 0.01 and 0.00001. Enrichment analysis was done using a Fisher exact test based on the COGs, using Claude ai for the script (REF).

### Phenotypic assays

#### Production of virulence factors

Individual colonies (four biological replicates) were picked and stabbed into LB agar plate containing 1.5% skimmed milk (assessing proteolytic activity; (49)), Cas assay plates made with 10X chelated LB (23); 0.5 g/L yeast extract supplemented with 4g/L D-mannitol (YEM) and agar (assessing mucoidy; (50)) and Columbian horse blood agar plate (assessing hemolysis activity; (51)). Plates were incubated at 37°C for 3 days. Inhibition circles were measured and confirmed by statistical analyses.

#### Antimicrobial sensitivity assay

Individual colonies were resuspended in 0.9% NaCl solution to obtain a turbidity of 0.5-0.63 McFarland. Bacterial suspensions were then spread onto Muller Hinton II agar plates and incubated overnight at 37°C with 0.002-32 µg/mL Meropenem (MEM) ETEST (Biomérieux), 10µg Ceftazidime (CAZ) and 25µg Sulphamethoxazole-Trimethoprim (TMP-SMX) Antimicrobial Susceptibility discs (Oxoid). Antimicrobial susceptibility was determined by following the recommendation from EUCAST (v15; https://www.eucast.org/fileadmin/src/media/PDFs/EUCAST_files/Breakpoint_tables/v_15.0_Breakpoint_Tables.pdf). For CMVs from the same isolate showing considerable visual changes in antibiotic susceptibility, three biological replicates were conducted on independent days, and the average of inhibition data were used to determine their susceptibility characteristic.

### Data availability

Genomes from this study are available in GenBank under the Bioproject: PRJNA1295108 (52, 53). Proteomic raw data deposited to the ProteomeXchange Consortium via the PRIDE partner repository with the dataset identifier PXD066041 (54).

## Results

### Isolation of TSV292_1 rough and TSV292_2 smooth colony morphotypes

In *Burkholderia,* CMV typically occurs upon removal of environmental pressure or during aging (13, 17). Because selective media for *Burkholderia pseudomallei* often contain antibiotics that can induce colony morphology changes (e.g., the wrinkled phenotype in *Bp*), the clinical isolate TSV292 was streaked directly onto LB agar supplemented with 0.1% congo red (CRLA) from a −80 °C glycerol stock to facilitate CMV distinction. On this medium, a mixed population was observed, consisting of flat colonies with a red center (type RI) and smoother, raised colonies (type SI) (**Fig. 1A**). The rough and smooth phenotypes were confirmed by isolating single colonies and subculturing them on CRLA or Ashdown agar (**Fig. 1B**). Both media yielded CMV types consistent with those previously reported for isolate TSV82 (23).

**Figure 1:**
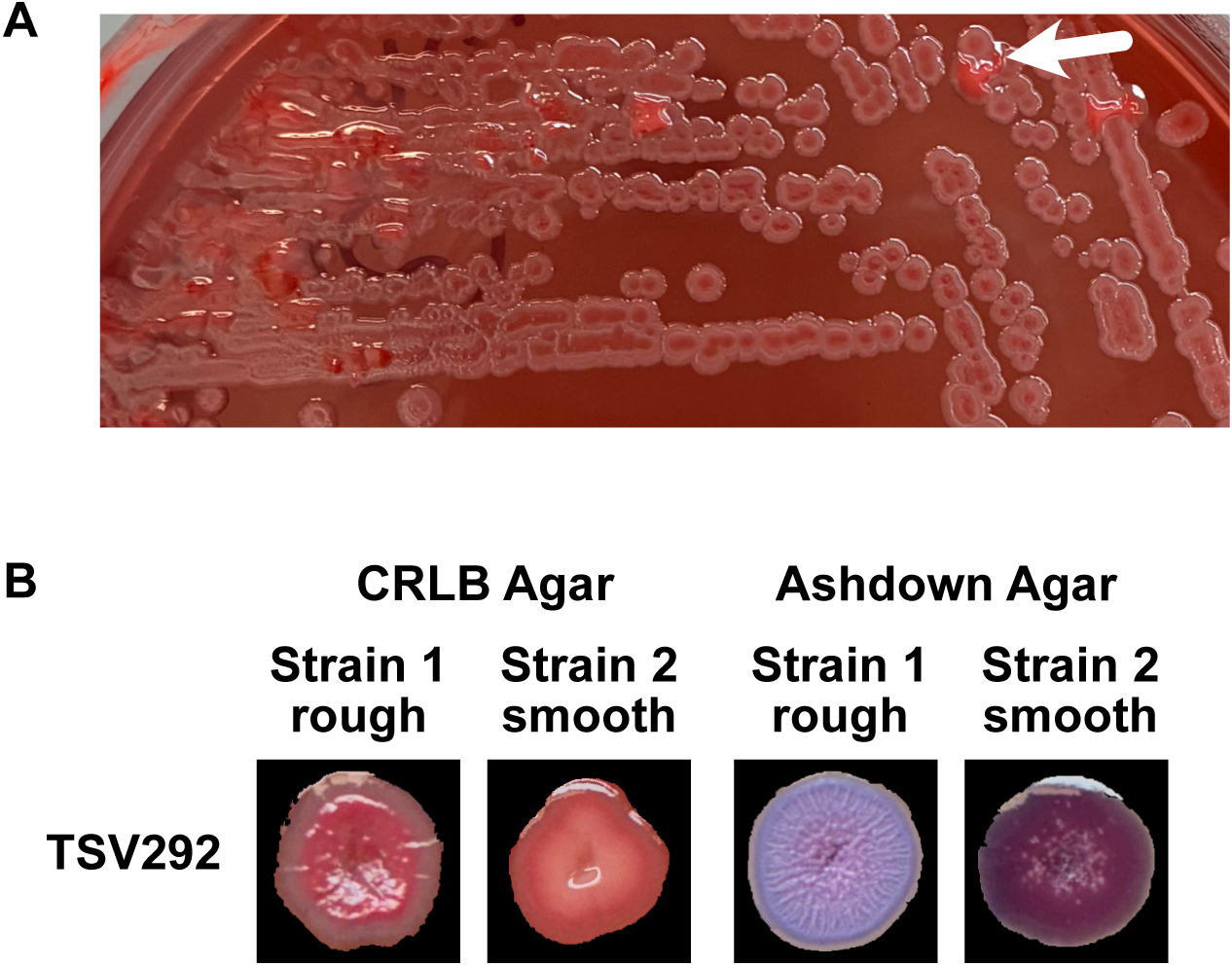
Colony morphologies of the two *Bp* strains isolated from TSV292. A) -80°C glycerol stock streak on CRLA, white arrow indicates the presence of second colony morphotype corresponding to TSV292_2 smooth strain. B) Colony morphology of both isolates after 3-days growth on both CRLA and Ashdown agar.

### Polyclonal strains masquerading as CMV in melioidosis infection

Genomic variation, master regulator expression or epigenetics are known drivers of CMV in *Burkholderia*; hence, we first investigated whether SNPs or insertion and deletions (indels) accounted for the distinct morphologies of TSV292. Illumina reads from TSV292 rough were mapped against the assembled genome of TSV292 smooth, using Snippy (33), which revealed 23,229 genomic modifications (**Table S1**). This unexpectedly high level of variation suggested that the two morphotypes were polyclonal rather than colony morphotype variants of the same isolate. Multi-locus sequence typing (MLST) further supported this conclusion: the rough isolate was identified as ST-2319 (TSV292_1; PubMLST sample ID:7631; previously reported by Gassiep and colleagues (24)), whereas the smooth isolate represented a novel sequence type (TSV292_2; PubMLST sample ID:7519; ST-2323) (**Table S2**). As simultaneous infection with multiple distinct *Bp* clones from the same patient is considered rare (25), we further investigated the genomic and proteomic profiles of these isolates to better understand their divergence.

### Comparative analysis of *Bp* ST-2319 and ST-2323

The average nucleotide identity (ANI) between the two genomes was 99.4%. TSV292_1 (rough) had a genome size of 7,262,464 bp with a GC content of 68.1%, while TSV292_2 (smooth) had a genome size of 7,352,845 bp with a GC content of 68.0%. Clustering analysis of whole-genome MLST (wgMLST) based on 6,259 allelic profiles of 1,225 available isolates in the PubMLST database (**Table S3**; as of August 2025) revealed two distinctly separate groups (**Fig. 2A**). Maximum-likelihood phylogenetic analysis showed that TSV292_1 rough clustered with *Bp* strains previously isolated from melioidosis patients admitted at Townsville University Hospital between 1997 and 2020 (24), whereas TSV292_2 smooth was more closely related to *Bp* strains isolated in Cambodia (**Fig. 2B**). TSV292_1 rough and TSV292_2 smooth shared >99.2% ANI with their closely related *Bp* isolates but also contain variable regions representing strain-specific regions (**Fig. S1**).

**Figure 2:**
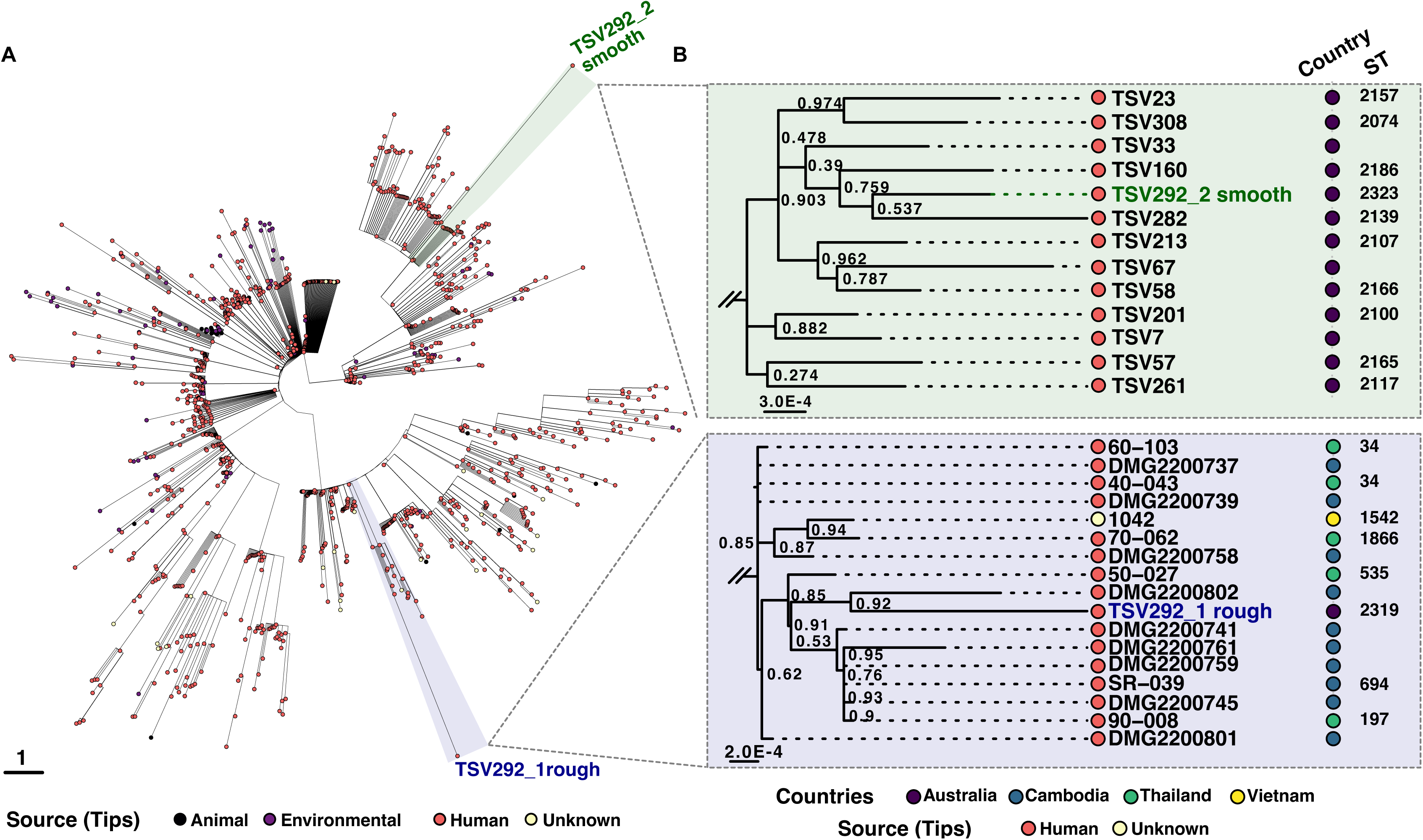
Two distinct *Bp* strains co-infecting the same patient suffering from melioidosis in 2017. A) Circular phylogenetic tree constructed using GrapeTree software showing relationships between isolates based on wgMLST allelic profiles. The tree represents a minimum spanning tree where branch lengths correspond to the number of allelic differences between isolates across genome loci. Each tip represents an individual isolate, with closer proximity indicating greater genetic similarity. Two reference strains are highlighted: TSV292_1 rough (blue) and TSV292_2 smooth (green). The scale bar represents 1 allelic difference. Isolates are colored by source as indicated in the legend. The analysis included 1,225 isolates characterized using 6,259 loci from the wgMLST scheme. B) Zoomed on closest isolated for both TSV292_1 rough (blue, right top panel) and TSV292_2 smooth (green, right bottom panel), using maximum likelihood phylogenetic analysis, using Fasttree, based on closest relative genomes.

Whole-genome alignment of the TSV292 co-isolates, using MAUVE algorithm (40), revealed synteny across the locally co-linear blocks and highlight genomic rearrangements likely as a result of recombination events in both chromosomes (**Fig. S2**). Alignment using the LASTZ tool (41, 42), further revealed divergence between TSV292_1 rough and TSV292_2 smooth in several regions (**Fig. S3–4**).

In TSV292_1 rough, regions absent in TSV292_2 smooth included: (i) a 7,465 bp segment containing a type II restriction enzyme and DNA Methyltransferase (DNA MTase) predicted to be responsible for m5C methylation on GTCGAC pattern (TSV292_R_RS_00500-00501) surrounded by IS3 family transposase; (ii) a 36,612 bp region including predicted phage elements and a singleton DNA MTase (TSV292_R_RS_02319); and (iii) a 61,103 bp region with 26 putative transposases consistent with lateral gene transfer, that also encoded the *Bp* adhesion (*bpa*) gene cluster—implicated in bacterial attachment and immune evasion (55) — suggesting functional differences in host interaction and pathogenic potential.

On chromosome 2 TSV292_1 rough carried a 43,430 bp region with an integrase (TSV292_R_RS_04679) and phages elements; a 36,084 bp region rich in transposases and viral genes, including a type III restriction DNA methyltransferase (TSV292_R_RS_03252; **Table S7**); a 21,283 bp region containing a putative origin of replication with a PAAR gene and a type III restriction DNA MTase (TSV292_R_RS_03525) suggestive of plasmid integration; and a 43,340 bp region containing phage elements (**Tables S4 and S5**).

By contrast, TSV292_2 smooth lacked multiple phage-associated and transposon-rich regions present in TSV292_1 rough, including loci encoding type II, III, and IV restriction MTases predicted to modify m4C, m5C, and m6A motifs (e.g., TSV292_RS_03205, TSV292_RS_03345, TSV292_RS_15530, TSV292_RS_15535, TSV292_RS_18580, TSV292_RS_26185; Tables S4, S6–S7). Differences also extended to the flagellar biosynthesis cluster, where the two genomes displayed altered gene order (Fig. S5).

Taken together, these results indicate that divergence between both strains involved acquisition or loss of phage elements, transposons, plasmid-associated regions, and DNA methylation systems. Such changes not only underscore the genome plasticity of *Bp* but also suggest functional consequences for immune evasion, horizontal gene transfer, and regulation of surface structures critical for infection.

### Distinct phenotypes in antibiotic sensitivity, mucoidy, hemolysis, siderophore production and proteolytic activity

To confirm the proteomic analyses, phenotypic assays assessing virulence-related traits, including antibiotic susceptibility were performed (**Table 1**), revealing notable differences between the two isolates. TSV292_2 smooth exhibited a higher resistance to trimethoprim-sulfamethoxazole (TMP-SMX), with no apparent difference in predicted antibiotic resistance by ARDaP (**Table S8-9**; (39)), and displayed increased mucoidy, consistent with higher exopolysaccharide (EPS) production when grown on YEM agar. In contrast to TSV292_1 rough, the TSV292_2 smooth strain lost its ability to lyse red blood cells and lacked proteolytic activity. Interestingly, both isolates produced a smooth-edged sector around the main colony when grown on Ashdown and blood agar, suggesting an active role in CMV emergence (**Fig. 3**).

**Figure 3:**
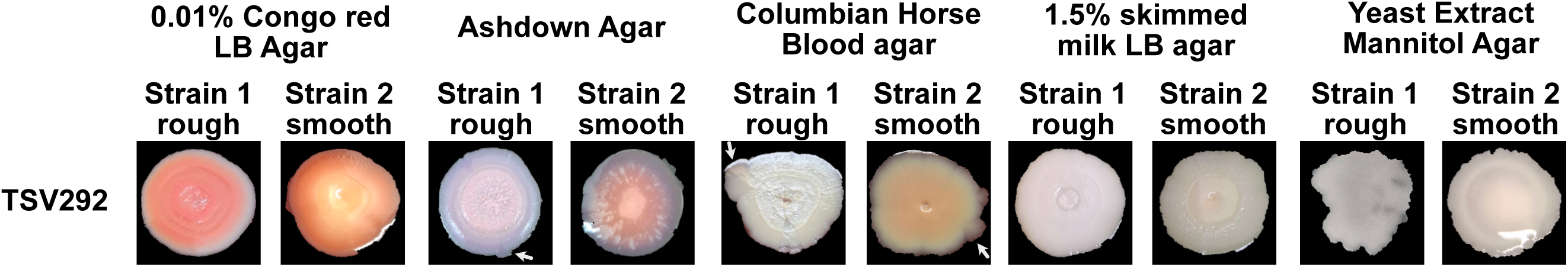
CMV emergence from both isolates occur after 8 days of growth on various media. White arrows indicate variants emergence from the main colony when culture on different media to study bacterial phenotypes.

**Figure 4:**
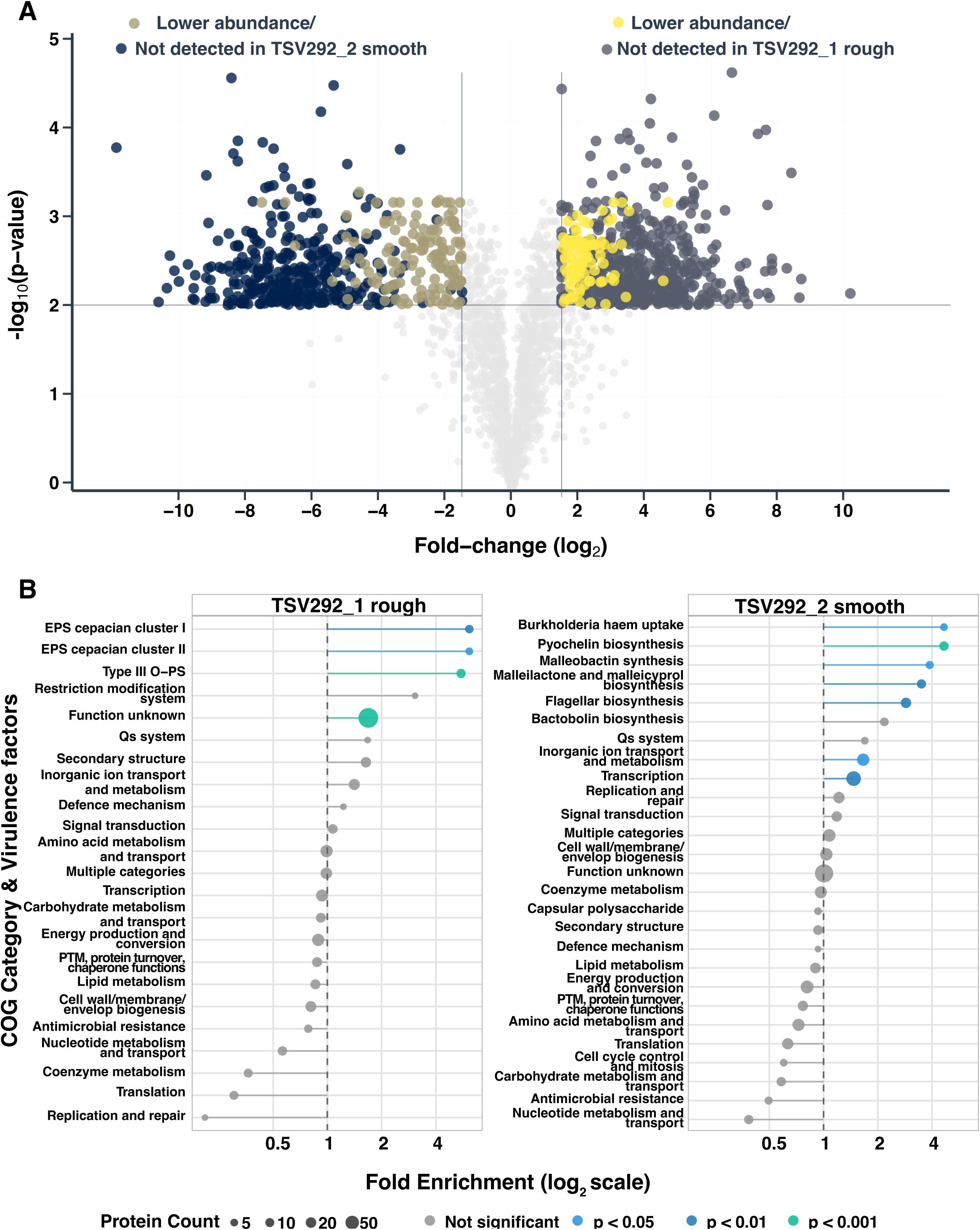
Proteomic profiling of both TSV292_1 rough and TSV292_2 smooth isolates. A) Volcano plot representing the significant abundance changes in proteins between both isolates. B) Enrichment analysis of virulence factors and COGs reflecting the proteins changes.

**Table 1.**
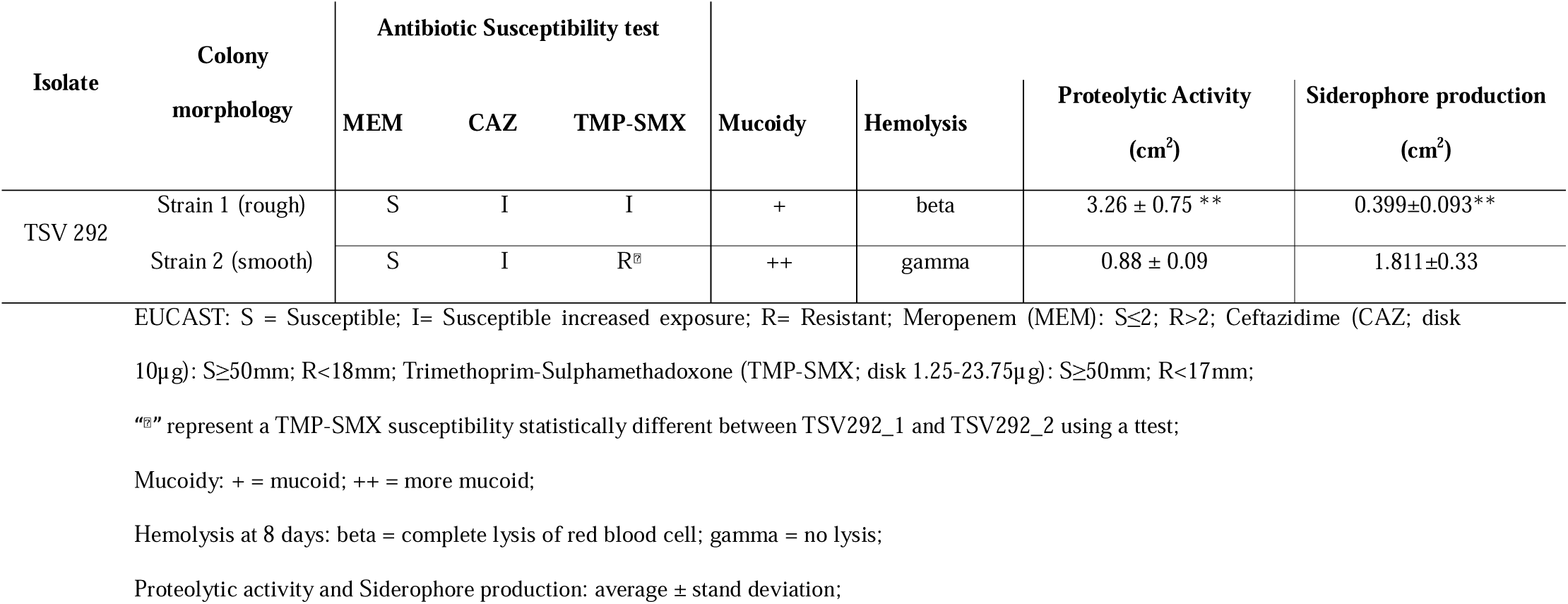
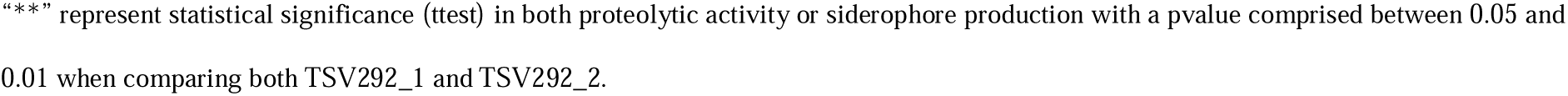
Phenotypic profile of both *Bp* TSV292_1 rough and TSV292_2 smooth.

### Proteomics confirms profile respectively belonging to rough and smooth *Bp* CMV morphotypes despite being two isolates

Proteomic profiling confirmed that the two isolates correspond to the expected profiles of rough and smooth *Bp* morphotypes, with differences in virulence- and resistance-associated pathways. In total, 1,015 proteins were differentially produced between the two isolates (**Fig. 3A; Table S10–11**).

In the TSV292_2 smooth strain, proteins associated with antimicrobial gene cluster biosynthesis were more abundant or exclusively detected. These included those involved in the production of pyochelin, malleobactin, malleilactone, bactobolin, and syrbactins (56–58). Notably SyrC/GlbC, part of the syrbactin cluster, is also known to influence *Bp* survival and replication within immune cells (56, 59). Additionally, proteins related to flagellar biosynthesis, the quorum sensing system 2 (BpsR2; belonging to the bactobolin biosynthesis cluster), and the Hmq system, which produces HMAQs with antimicrobial activity and known interaction with QS systems, were more abundant or exclusively detected (58, 60, 61). In contrast, the TSV292 rough morphotype showed higher abundance or exclusive detection of proteins involved in exopolysaccharide (EPS) production, type III O-polysaccharide (OPS) synthesis, DNA methyltransferase (MTase) responsible for the CACAG methylation motif, the quorum sensing system 3 (BpsI3/BpsR3), and the master regulator ScmR (**Fig. 3B**).

Proteins from antibiotic resistance genes (11, 39, 62, 63), predicted using the CARD database (**Table S12;** (38)), were generally not detected or detected at lower abundance in TSV292_2 smooth compared with the rough isolate (**Table 2**). This includes both AmrR, the TetR family transcriptional regulator that controls the expression of the AmrAB-OprA efflux pump, and BpeT, the transcriptional regulator of the BpeEF-OprC efflux pump, potentially explaining the observed TMP-SMX resistance in the smooth strain in Table 1 (39).

**Table 2:**
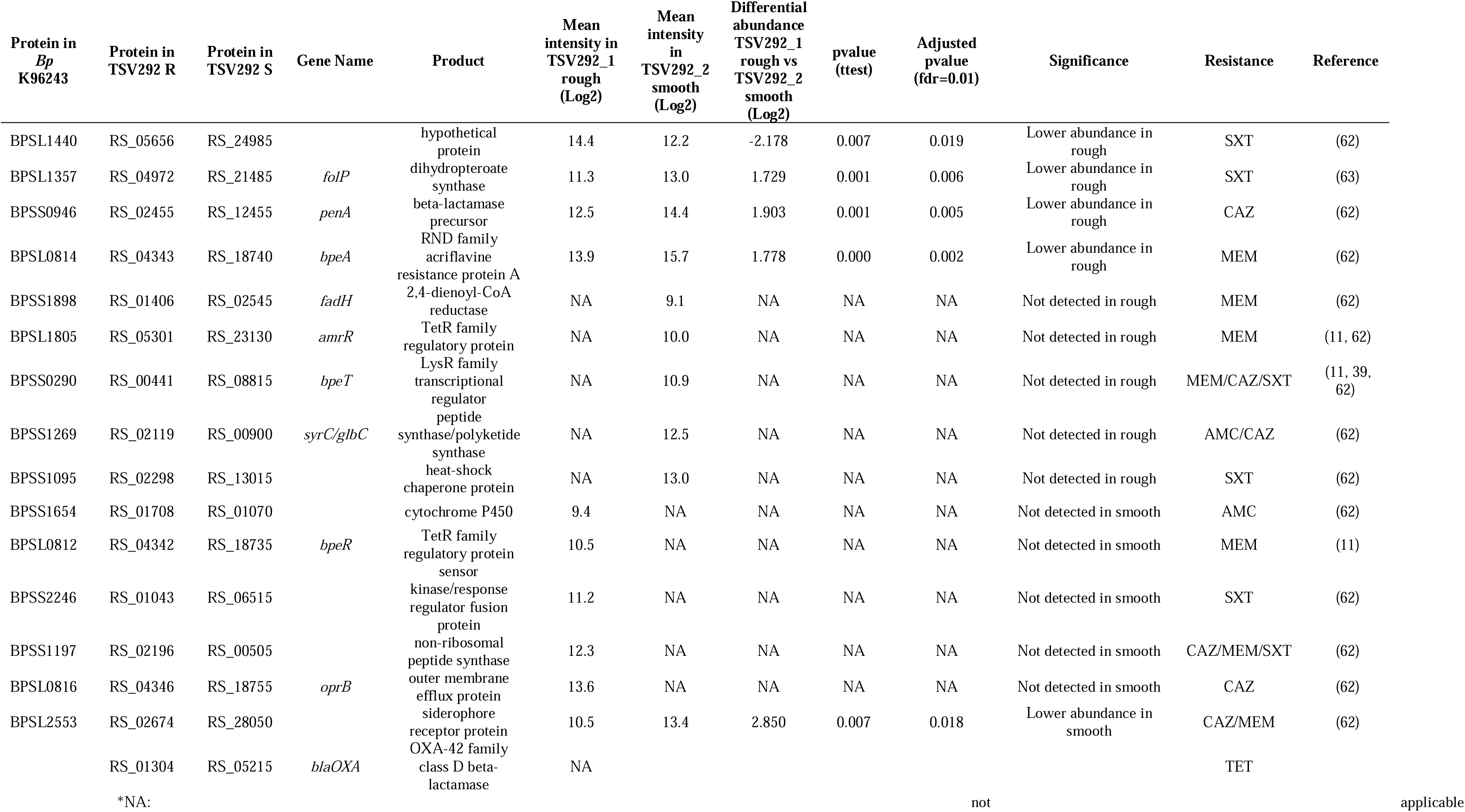
Significance differences in the abundance of protein involved in antibiotic resistance.

Together these results indicate a difference in adaptations between the two strains, with TSV292_2 smooth adopting an environmental-like smooth form and TSV292_1 rough displaying a pathogenic rough form (15, 58).

### Methylome showed no overall difference in methylated sites between TSV292 rough and smooth

Predicted restriction enzyme methylated motifs were consistent with the one described by Nandi et al. (64) (**Table S7&13**). Given the observed difference in abundance of the DNA MTase responsible for the CACAG methylation motif (TSV292_R_RS_03660/03661 in rough colony morphotype TSV292_RS_15080/15085 in the smooth, for which proteins were not detected in TSV292_2 smooth while detected in TSV292_1 rough isolates), we further investigated the methylome profiles of both isolates (**Table S13**). Despite the presence of distinct abundance of DNA MTases, no significant differences were observed in methylated sites across the genomes of TSV292 rough and smooth for motifs CACAG, GTWWAC and CATCAG.

## Discussion

Melioidosis causes an estimated 179,000 cases and 90,000 deaths worldwide each year (4). As *Bp* is intrinsically resistant to antibiotics, treatment relies on last-resort agents. Emerging resistance remains a major concern (8–10). While simultaneous polyclonal *Bp* infections are considered rare (25), here we report the first such case in Queensland, isolated from a patient swab in 2018.

A key diagnostic challenge revealed by this study is that polyclonality can be mistaken for CMV. Current workflows, typically single-colony purification with selection of a typical wrinkled morphotype followed by MALDI-TOF or sequencing, cannot reliably distinguish between true CMV and mixed-strain infections. This diagnostic blind spot risks underestimating strain diversity and may lead to inappropriate clinical decisions and treatment failures (12). Because *Burkholderia* species exhibit extensive colony morphotype plasticity with a particular mucoid/wrinkled colony morphotype (13, 17, 19), relying on phenotypic assessment alone can mask genetically distinct pathogens. In this case, two morphotypes that appeared as rough and smooth variants on Congo red agar and mucoid/wrinkled on Ashdown selective media were in fact distinct strains.

We also investigated DNA methylation, given its known role in CMV and intraspecies recombination (65). While global methylation patterns appeared similar between the isolates, the absence of detectable DNA MTases by LC-MS/MS in the smooth morphotype mirrored our prior findings in smooths CMVs (23). This could reflect biological differences in MTase stability or abundance but may also stem from limitations of single-replicate methylome analysis in detecting subtle variation.

By integrating genomic, proteomic, and phenotypic analyses, we demonstrate how distinct co-infecting strains can mimic CMV, with smooth and rough colony morphotypes differing in the production of proteins involved in virulence, stress tolerance, drug responses and adaptation—features with direct implications for patient management. Importantly, outbreak tracing and phylogenomic inference are highly vulnerable to such hidden co-infection; clonal verification should therefore precede any evolutionary or epidemiological analyses of *Bp*. Our findings suggest that covert polyclonality, rather than ultrafast evolution, is a major driver of observed genomic diversity in this pathogen. High-quality, closed genomes from long-read assemblies, combined with read-level deconvolution using short-read sequencing, should be standard practice before calling variants.

Ultimately, this work highlights the urgent need for strain-level diagnostic approaches that integrate multi-omics and phenotypic data. Building comprehensive reference databases of such profiles will enable predictive tools to guide diagnosis, treatment, and surveillance. These advances will not only improve patient outcomes but also yield more accurate epidemiological insights, deepening our understanding of *Bp* biology.

## Acknowledgment

This work was supported by AIMI and UTS Faculty of Science seed funds to Dr Pauline M.L. Coulon, UTS Strategic Research Accelerator funding (SRA 2726229) to Prof Garry S.A. Myers and EL2 Investigator grant from the NHMRC (APP2033851) to Dr Patrick N.A. Harris. Experimental work was done at UQCCR which would not have been possible without the ISME scholar funding awarded to Dr Pauline M.L. Coulon.

P.M.L.C: Concept and visualization, Funding, Experimental work; K.M: Experimental work and reviewing, P.R.C: Genomic comparison; M.P and J.T: DNA libraries and sequencing; A.L, E.R and S.R: sample preparation and LCMS runs for the proteomic experiment; I.G & P.N.A.H: Reviewing and Funding; G.S.A.M: Reviewing and Funding.

