## supplementary figures for "Co-isolation of genetically distinct *Burkholderia pseudomallei* strains from a single patient in North Queensland"

Pauline M.L. Coulon^1^, Piklu Roy Chowdhury^1^, Aven Lee^2^, Edita Ritmejeryte^2^, Miranda Pitt^1^, Joyce To^1^, Kay Ramsay^3^, Sarah Reed^2^, Patrick N. A. Harris^3^ and Garry S.A. Myers^1^

Affiliations:

^1^Australian Institute for Microbiology and Infection, Faculty of Science, University of Technology Sydney, NSW, Australia

^2^ Mass Spectrometry Facility, University of Queensland, Centre for Clinical Research, Queensland, Australia

^3^The University of Queensland Centre for Clinical Research (UQCCR), Faculty of Medicine, The University of Queensland, Queensland, Australia

Corresponding author:

Pauline M.L. Coulon, Australian Institute for Microbiology and Infection, Faculty of Science, University of Technology Sydney, NSW, Australia, +61390353555,

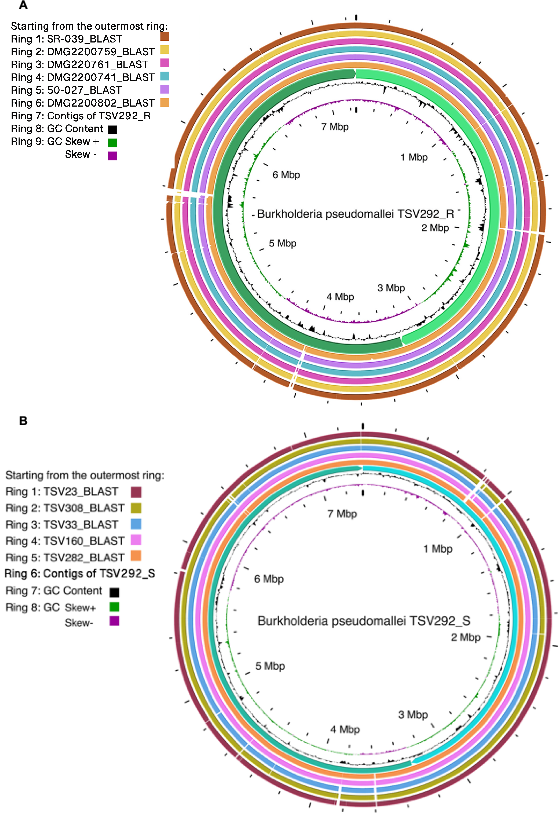

**Figure S1**: **Genome alignment of TSV292_1 (rough) and TSV292_2 (smooth) with their respective closest isolates genomes.**

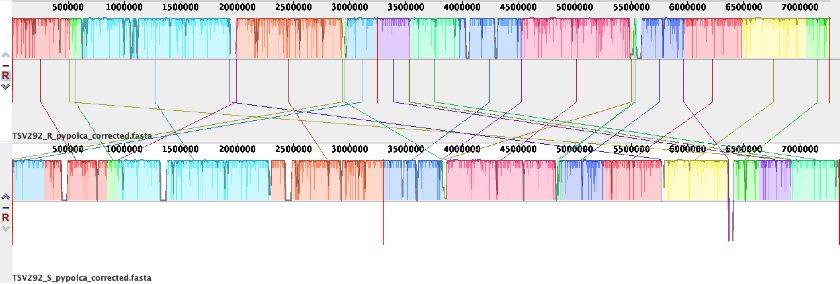

**Figure S2: MAUVE alignment of both** **TSV292_1 (rough) and TSV292_2 (smooth) against each other.** The red line delimits both chromosomes in each genome. Contig 1 is one the left and contig 2 is on the right. Each color correspond to an identical region in both genome.

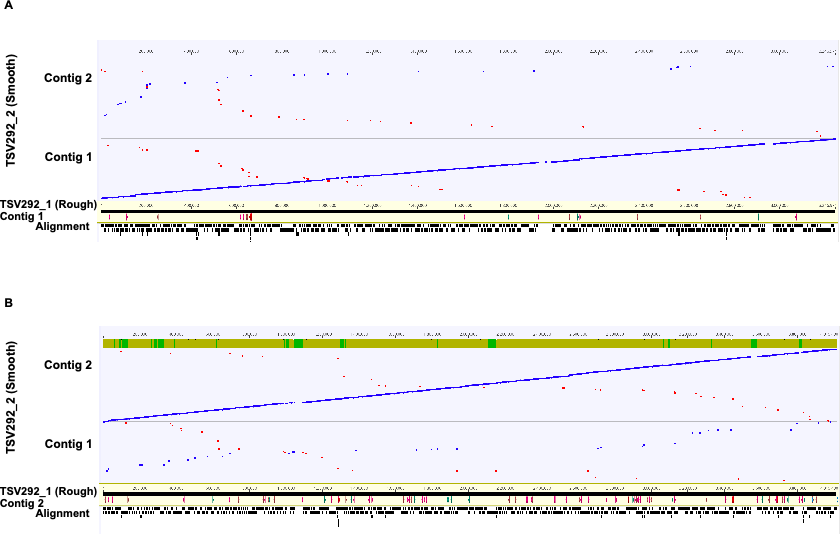

**Figure S3**: **Genome alignment of TSV292_2 (smooth) on TSV292_1 (rough) using LASTZ alignment tool.** Forward alignments between the reference TSV292_1 (rough) contig 1 (**A**) and contig 2 (**B**) (x-axis) and TSV292_2 (smooth) query (y-axis) genomes are shown in blue, indicating regions of conserved synteny in the same orientation. Alignments in red represent sequences aligned in the reverse-complement orientation, corresponding to inverted segments relative to the reference. Gaps with no alignment indicate regions unique to one genome, missing from the other, or below the similarity threshold for alignment.

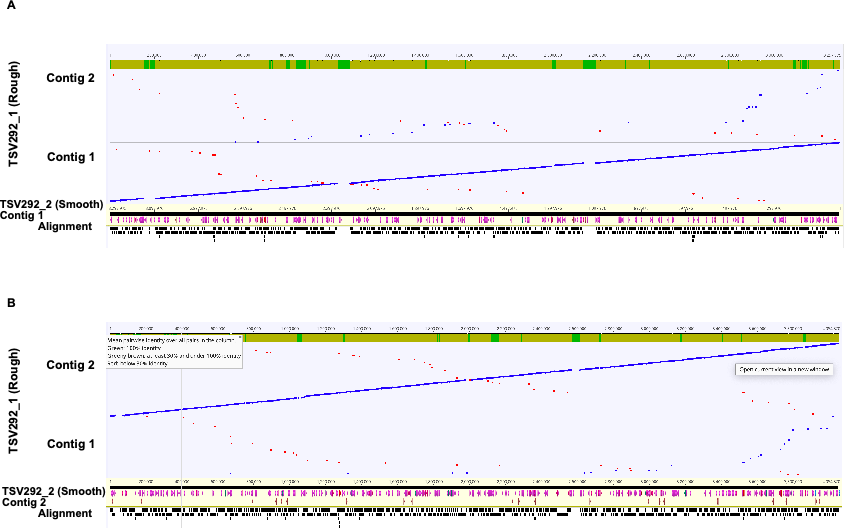
**Figure S4**: **Genome alignment of TSV292_1 (rough) on TSV292_2 (smooth) using LASTZ alignment tool.** Forward alignments between the reference TSV292_2 (smooth) contig 1 (**A**) and contig 2 (**B**) (x-axis) and TSV292_1 (rough) query (y-axis) genomes are shown in blue, indicating regions of conserved synteny in the same orientation. Alignments in red represent sequences aligned in the reverse-complement orientation, corresponding to inverted segments relative to the reference. Gaps with no alignment indicate regions unique to one genome, missing from the other, or below the similarity threshold for alignment.

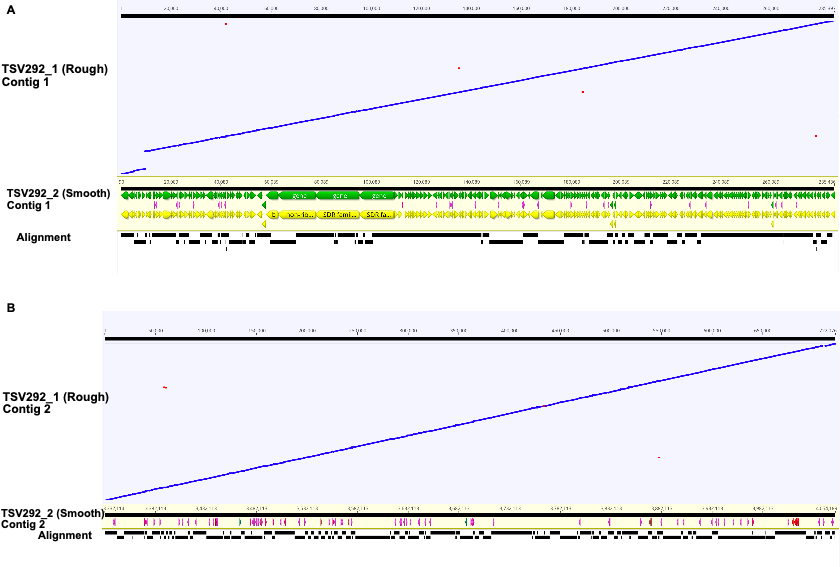

**Figure S4**: **Genome alignment of TSV292_1 (rough) on TSV292_2 (smooth) using LASTZ alignment tool of the missing dot plot of Figure S2-3.** Forward alignments between the reference TSV292_2 (smooth) of the missing part in figure S3 for contig 1 (**A**) and contig 2 (**B**) (x-axis) and TSV292_1 (rough) query. This figure demonstrated that there are no missing genomic regions of 285,397bp and 722,076 bp in contig 1 and 2 of respectively.

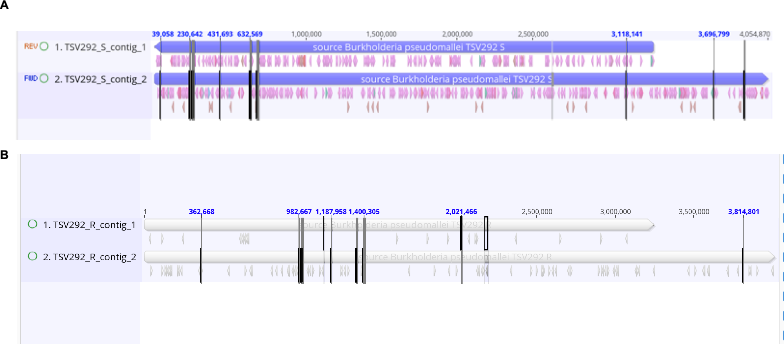

**Figure S5: Highlight of the order of genes with an identified as flagellar product.**

A) TSV292_2 (smooth) genome. B) TSV292_1 (rough) genome.
